# Notch signaling regulates immunosuppressive tumor-associated macrophage function in pancreatic cancer

**DOI:** 10.1101/2023.01.11.523584

**Authors:** Wei Yan, Nina G. Steele, Samantha B. Kemp, Rosa E. Menjivar, Wenting Du, Eileen S. Carpenter, Katelyn L. Donahue, Kristee L. Brown, Valerie Irizarry-Negron, Sion Yang, William R. Burns, Yaqing Zhang, Marina Pasca di Magliano, Filip Bednar

**Author notes:** Henry Ford Hospital, Detroit, MI 48202, USA. Department of Surgery, Johns Hopkins University School of Medicine, Baltimore, MD 21287, USA. Corresponding authors. **Corresponding authors:** Filip Bednar (FB), Mail: 2210 Taubman Center, 1500 E Medical Center Dr. Ann Arbor, MI 48109, Marina Pasca di Magliano (MPdM), Mail: Rogel Cancer Center Rm. 6306, 1500 E Medical Center Dr. Ann Arbor, MI 48109, Yaqing Zhang (YZ), Mail: Rogel Cancer Center Rm. 6110, 1500 E Medical Center Dr. Ann Arbor, MI 48109. **Author contributions**, FB and MPM directed the study. WY, YZ, FB and MPM designed experiments. KLD, KLB, WD, REM, ELLO, JL, WY, SBK, MB, FL, CAL, HCH, TLF performed the experiments and generated data. FL, CAL, HCH, TLF analyzed and interpreted data. WY, YZ and FB wrote the manuscript, and FB and MPM edited and all authors approved the final version.

## Abstract

Pancreatic ductal adenocarcinoma (PDA) continues to have a dismal prognosis. The poor survival of patients with PDA has been attributed to a high rate of early metastasis and low efficacy of current therapies, which partly result from its complex immunosuppressive tumor microenvironment. Previous studies from our group and others have shown that tumor-associated macrophages (TAMs) are instrumental in maintaining immunosuppression in PDA. Here, we explored the role of Notch signaling, a key regulator of immune response, within the PDA microenvironment. We identified Notch pathway components in multiple immune cell types within human and mouse pancreatic cancer. TAMs, the most abundant immune cell population in the tumor microenvironment, express high levels of Notch receptors with cognate ligands such as *JAG1* expressed on tumor epithelial cells, endothelial cells and fibroblasts. TAMs with activated Notch signaling expressed higher levels of immunosuppressive mediators including arginase 1 (*Arg1*) suggesting that Notch signaling plays a role in macrophage polarization within the PDA microenvironment. Combination of Notch inhibition with PD-1 blockade resulted in increased cytotoxic T cell infiltration, tumor cell apoptosis, and smaller tumor size. Our work implicates macrophage Notch signaling in the establishment of immunosuppression and indicates that targeting the Notch pathway may improve the efficacy of immune-based therapies in PDA patients.

## Introduction

90% of pancreatic ductal adenocarcinoma (PDA) patients die within five years of their diagnosis (1). A key reason for this poor prognosis is resistance to existing therapies. This resistance is partly regulated by oncogenic KRAS signaling within the pancreatic tumor epithelium (2). Genetically engineered mouse models expressing oncogenic KRAS in the pancreas recapitulate the histologic progression of human PDA with formation of preneoplastic lesions including acinar-ductal metaplasia (ADM), Pancreatic Intraepithelial Neoplasia (PanIN), and pancreatic adenocarcinomas with metastases (3–6). Based on the observations from these models and human patients, it has become clear that epithelial oncogenic KRAS signaling promotes the formation of a complex immunosuppressive fibroinflammatory stroma, which also contributes to treatment resistance (7).

Notch signaling is one of the core pathways dysregulated in PDA downstream of oncogenic KRAS (8). The core components of Notch signaling in mammals include the Notch transmembrane receptors - Notch1-4 - and five membrane-bound ligands - Jagged1 and 2 and Delta-like ligand 1, 3, and 4 (9). Cell-cell contact and ligand–receptor interaction between neighboring cells lead to a series of proteolytic events culminating in the γ-secretase-mediated cleavage of the intracellular Notch domain (NICD), which is then released from the plasma membrane and translocates into the nucleus where it binds to the DNA-binding protein CBF1/RBPJκ. This complex also recruits the transcriptional co-activator mastermind-like (MAML) and leads to transcriptional activation of Notch target genes including the Hes family of transcription factors (9). Notch epithelial signaling regulates pancreatic neoplastic progression in genetically engineered mouse models (10–14) and inhibitors targeting γ-secretase activating Notch proteolytic cascade have been developed (15). Unfortunately, clinical trials based on these approaches have so far not yielded improved survival (16,17). One reason for this lack of efficacy is incomplete understanding of how Notch regulates the PDA tumor microenvironment.

Notch pathway regulates multiple aspects of the tumor immune response including T cell differentiation and maturation and myeloid compartment functionality (18–20). In the context of the PDA TME, Notch inhibition via γ-secretase inhibition (GSI) led to increased intratumoral hypoxia and sensitivity of tumor epithelium to systemic therapy (21). In addition, Notch activation in vitro has been implicated in promoting M1-like, anti-tumor macrophage polarization (22,23). Genetic approaches to activate or inhibit Notch signaling and transcriptional response in the myeloid compartment of an autochthonous mouse model of pancreatic cancer demonstrated increased productive cytotoxic T cell and macrophage anti-tumor response associated with Notch stimulation (23). However, contrary to this observation, breast cancer models with Notch activation within tumor-associated macrophages (TAMs) demonstrated protumorigenic, M2-like TAM polarization, T cell inhibition, and a blunted anti-tumor immune response (24). The state of Notch signaling in the myeloid compartment of human PDA patients remains unclear. In our work, we address the activation state of Notch within multiple compartments of primary human and mouse PDA and how the myeloid compartment and tumor immune response is shaped in the presence of pharmacological Notch and immune checkpoint inhibition.

## Materials and Methods

### Study design

The objective of this project was to study the role of Notch signaling in TME on shaping the functional fate of tumor-associated macrophages of in male and female mice bearing pancreatic tumors and to provide insight into the role of Notch pathway in regulating immune treatment.

### Mice

CBF:H2B-Venus mice (25) were gifts from Dr. Sunny Wang, University of Michigan. By using multiple CBF1 binding sites together with a subcellular-localized, genetically-encoded fluorescent protein, H2B-Venus, the CBF:H2B-Venus transgenic strain of mice is capable of faithfully recapitulating Notch signaling at single-cell resolution (5). Venus mice were generated by crossing with C57BL/6 mice. Male and female mice were included equally. All animal studies were conducted in compliance with the guidelines of Institutional Committees on Use and Care of Animals at the University of Michigan.

### Animal experiments

To establish the orthotopic pancreatic cancer model, 5×10^4^ of 7940B cells (32) and 1X10^5^ mT3-2D cells (33), both derived from KPC mouse tumors (Pdx1-Cre; LSL-Kras^G12D/+^; LSL-Trp53^R172H/+^) in C57BL/6J background, were injected into Venus mice. Cells were tested for mycoplasma free by MycoAlertTM PLUS Mycoplasma Detection Kit (Lonza) and passage 15-20 were used for all experiments. γ-secretase inhibitor (GSI) Crenigacestat (LY3039478, Selleckchem Chemicals, Houston, TX) was given at 8 mg/kg by oral gavage every another day for Notch inhibition. Anti-PD1 treatment: Purified anti-mPD-1 antibody (BioXcell #BE0033-2; clone J43) was used for in vivo PD-1 blockade at a dosage of 200 μg/mouse through i.p. injection, repeated twice per week.

### Cell culture

All cells were cultured in IMDM supplemented with 10% FBS and 1% penicillin/streptomycin (Gibco). Mouse pancreatic cancer cell line 7940B was used to generate conditioned medium (CM). CM was filtered through 0.2 μm filter before use. For in vitro tumor-associated macrophage polarization, bone marrow derived myeloid cells (BMDM) were treated with CM (CM diluted 1:1 in fresh IMDM with 10% FBS) for 7 days for macrophage polarization; In addition, BMDM co-cultured with 7940B cells was used to polarize differentiated macrophages as previously described (34).

### Histopathological analysis

Hematoxylin and eosin (H&E), immunohistochemical and immunofluorescent staining were performed on formalin-fixed, paraffin embedded mouse pancreatic tissues as described before (Zhang et al., 2013a). Antibodies used are listed in Supplementary file 1. For immunofluorescence, Alexa Fluor (Invitrogen) secondary antibodies were used. Cell nuclei were counterstained with Prolong Gold with DAPI (Invitrogen). Images were taken with Olympus BX-51 microscope, Olympus DP71 digital camera, and DP Controller software. The immunofluorescent images were acquired using the Olympus IX-71 confocal microscope and FluoView FV500/IX software.

### Flow cytometric analysis and sorting

Single-cell suspensions of fresh spleen or pancreas were prepared as previously described (Zhang et al., 2013b) and stained with fluorescently conjugated antibodies listed in Supplementary file 1. Flow cytometric analysis was performed on a Cyan ADP analyzer (Beckman Coulter) and data were analyzed with Summit 4.3 software. Cell sorting was performed using a MoFlo Astrio (Beckman Coulter). Myeloid cells (DAPI-EGFP-CD45+CD11b+), epithelial cells (DAPI-EGFP+CD45-) and fibroblasts (DAPI-EGFP-CD45-CD11b-CD31-CD3-) were sorted and lysed in RLT buffer (Qiagen). Total RNA was prepared using RNeasy (Qiagen) and reverse-transcripted using High Capacity cDNA Reverse Transcription kit (Applied Biosystems).

### Quantitative RT-PCR

Samples for quantitative PCR were prepared with 1X SYBR Green PCR Master Mix (Applied Biosystems) and various primers (primer sequences are listed in Supplementary file 2). All primers were optimized for amplification under reaction conditions as follows: 95°C 10mins, followed by 40 cycles of 95°C 15 secs and 60°C 1 min. Melt curve analysis was performed for all samples after the completion of the amplification protocol. Cyclophilin A was used as the housekeeping gene expression control.

### Single-cell RNA sequencing

Single-cell suspensions of pancreatic tumors were derived as previously described (35). Dead cells were removed using MACS^®^ Dead Cell Removal Kit (Miltenyi Biotec Inc.). Single-cell cDNA library was prepared and sequenced at the University of Michigan Sequencing Core using the 10x Genomics. Samples were run using paired end 50 cycle reads on HiSeq 4000 (Illumina) or the NovaSeq 6000 (Illumina) to the depth of 100,000 reads per cell. The raw data were aligned to either mm10 or hg19 for mouse and human, respectively, then data were filtered using Cellranger count V3.0.0 with default settings at the University of Michigan, Advanced Genomics Core. R Studio V3.5.1 and R package Seurat version V3.0 was used for downstream single cell RNA-seq data analysis similarly as previous described (24,36). Data were initially filtered to only include all cells with at least 200 genes and all genes in greater than 3 cells. Data were initially normalized using the NormalizeData function with a scale factor of 10,000 and the LogNormalize normalization method. Variable genes were identified using the FindVariableFeatures function. Data were assigned a cell cycle score using the CellCycleScoring function and a cell cycle difference was calculated by subtracting the S phase score from the G2M score. Data were scaled, centered and batch corrected using linear regression on the counts, the cell cycle score difference and run ID using the ScaleData function. Principal Component Analysis (PCA) was run with the RunPCA function using the previously defined variable genes. Violin plots were then used to filter data according to user-defined criteria. Cell clusters were identified via the FindClusters function. FindMarkers table was created, and clusters were defined by user-defined criteria. Raw KPC mouse data are available at the NCBI’s Gene Expression Omnibus database (GSM6127792) and were originally published in (26). Human scRNA-seq data were previously published (24). Raw human data are available at the National Institutes of Health (NIH) dbGaP database (phs002071.v1.p1) and processed data are available at NIH Gene Expression Omnibus (GEO) database (GSE155698).

### Statistical analysis

Graphpad Prism six software was used for all statistical analysis. All data were presented as means ± standard error (SEM). Intergroup comparisons were performed using Two-tailed unpaired t-test, and p<0.05 was considered statistically significant.

### Study approval

All animal studies were conducted in compliance with the guidelines of the Institutional Animal Care & Use Committee (IACUC) at the University of Michigan. Patient selection/sample procurement: patients over the age of 18 referred for diagnostic endoscopic ultrasound of a pancreas mass lesion suspected of PDAC were consented according to IRB HUM00041280. Up to 2 extra passes were taken for research after biopsy obtained for clinical use. Surgical specimens were obtained from patients referred for Whipple or distal pancreatectomy according to IRB HUM000025339. Written informed consent forms were obtained from the patients, and the studies were conducted in accordance with recognized ethical guidelines. Human patient studies were approved by Institutional Review Boards of the University of Michigan Medical School.

### Data availability statement

All sequencing data used within this manuscript is publicly available on the Gene Expression Omnibus - GSM6127792 and GSE155698 – and in the NIH database of Genotypes and Phenotypes (dbGaP) for the raw human sample sequences - phs002071.v1.p1.

## Results

### Single-cell RNA sequencing reveals Notch pathway component expression in the human and mouse pancreatic tumor microenvironment

To explore Notch signaling pathway component expression in the human PDA microenvironment, we analyzed our previously published single cell RNA sequencing (RNA-seq) dataset of 16 human pancreatic cancer samples as well as 3 adjacent benign or normal pancreata (25). In this dataset, we captured 13 different cell populations in both PDA and adjacent/normal pancreata (Figure S1A). Gene expression profiling identified Notch signaling pathway components across all cell clusters at various levels except for acinar cells. In general, Notch signaling pathway gene expression was higher in PDA compared to adjacent/normal pancreata (Figure S1A). In particular, Notch receptors were highly expressed in cancer epithelial cells, endothelial cells, fibroblasts and subsets of immune cells including multiple myeloid populations (Figure S1A). The Notch pathway ligands such as *JAG1* were mainly expressed by epithelial cells, endothelial cells and fibroblasts (Figure S1A). Notch pathway canonical target gene *HES1* expression was enriched in epithelial cells, endothelial cells, fibroblasts, myeloid cells, and mast cells (Figure S1A). Co-immunofluorescent staining confirmed expression of HES1 in fibroblasts and macrophages in human PDA microenvironments (Figure S1B).

To further map the Notch signaling pathway within the immune system, we focused on gene profiling in immune cells only. We clustered the immune cells in 9 clusters (Figure 1A and S1C), containing multiple myeloid cell populations. Among those, monocytes, granulocytes and macrophages had high level of Notch receptors - primarily *NOTCH 1* and *2* (Figure 1B). The Notch target *HES1* was expressed by mast cells and monocyte and macrophage populations at relatively high levels compared to other immune cells (Figure 1B-C). In addition, *HES1* expression was upregulated in immune cells from pancreatic tumors compared to those from adjacent/normal pancreata, indicating that activation of Notch signaling in the myeloid compartment may affect tumor microenvironment regulation in pancreatic cancer (Figure 1D). The myeloid compartment also contained high expression levels of the proteolytic enzymes (*ADAM10*, *ADAM17*) involved in the Notch proteolytic cascade as well as downstream Notch signaling components (*MAML1/2*, *RBPJ*) suggesting the presence of a fully reconstituted Notch pathway, which may play a role in shaping the myeloid compartment within the PDA microenvironment.

**Figure 1.**
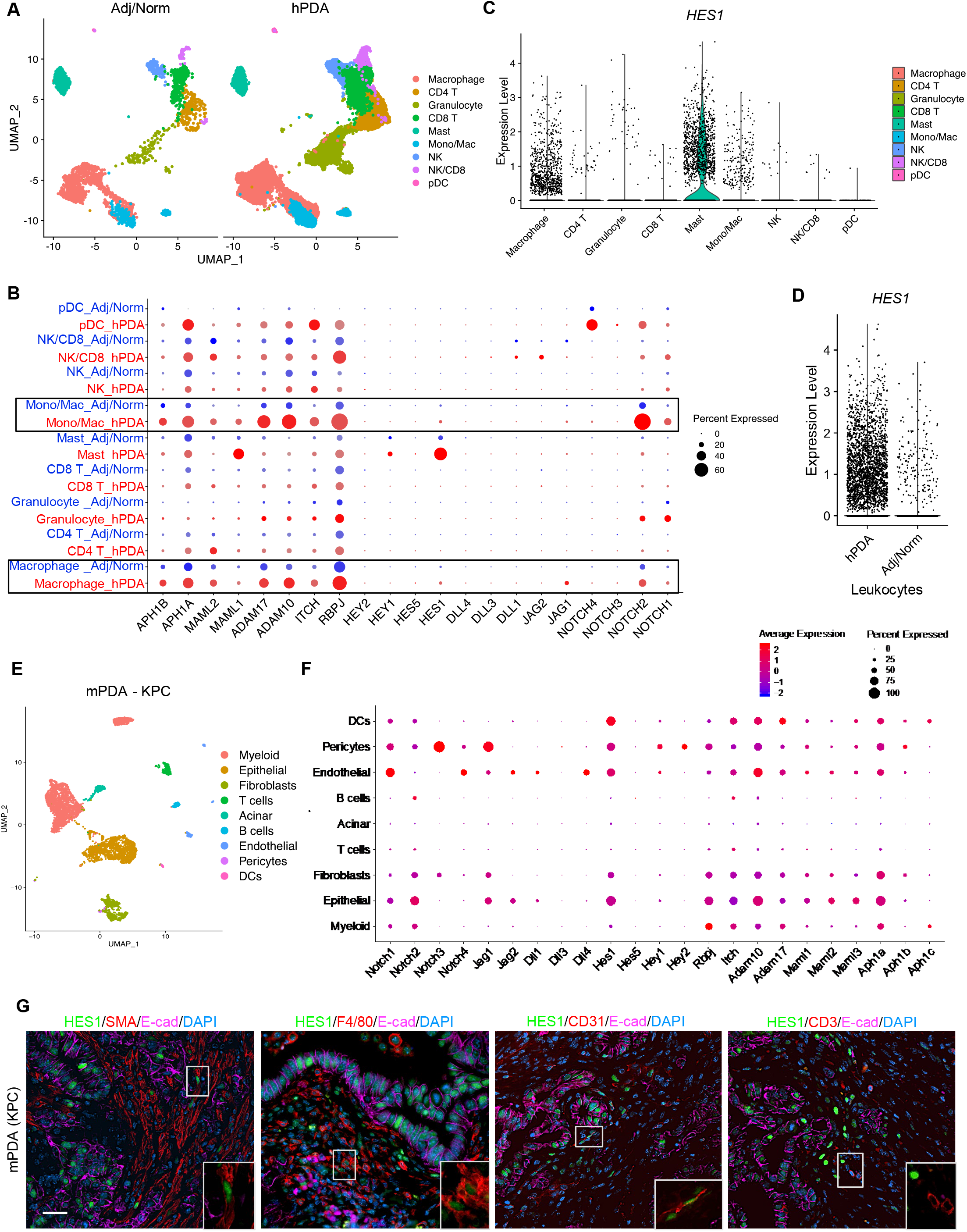
Notch activation in the tumor microenvironment of PDA. (A) UMAP plot of immune cells identified from single-cell RNA sequencing analysis with human pancreatic cancer samples (n=16) and adjacent benign/normal tissues (n=3), color-coded by their associated cluster. (B) Dot plot showing expression of Notch pathway genes across all immune cell clusters identified in the single-cell RNA sequencing analysis of human pancreatic samples. Size of dots represents percentage of cells expressing a particular gene and intensity of color indicates level of mean expression. (C) Violin plot of single cell RNA sequencing analysis showing expression level of *HES1* in different immune cell populations derived from human pancreatic cancer samples. (D) Violin plot of single cell RNA sequencing analysis comparing expression level of *HES1* in all leukocytes between human pancreatic cancer samples and adjacent benign/normal tissues. (E) UMAP plot of cell populations identified from single-cell RNA sequencing analysis with KPC mouse pancreatic cancer, color-coded by their associated cluster. (F) Dot plot showing Notch pathway genes across all clusters identified in the single-cell analysis of mouse KPC tumor. Size of dots represents percentage of cells expressing a particular gene and intensity of color indicates level of mean expression. (G) Co-immunofluorescent staining for HES1 (green), SMA, F4/80, CD31 or CD3 (red), E-cad (magenta) and DAPI (blue) in mouse KPC tumor sample. Scale bar 50 μm.

To address if the existing mouse models of pancreatic cancer recapitulate our observations from human tumors, we performed similar single-cell RNA-seq analysis within the Ptf1a^Cre/+^, Kras^LSL-G12D/+^, Tp53^LSL-R172H/+^ (KPC) mouse model (26) (Figure 1E). We observed high expression of Notch receptors, downstream signaling components, and *Hes1* in cancer epithelial cells, endothelial cells, pericytes, fibroblasts and myeloid cells including dendritic cells and macrophages (Figure 1F). Co-immunofluorescent staining confirmed abundant expression of HES1 in fibroblasts, macrophages, and endothelial cells in the KPC tumor microenvironment (Figure 1G). Our results support presence of active Notch signaling within multiple components of the PDA TME including macrophages.

### Macrophages in the pancreatic tumor microenvironment have active Notch signalings

To evaluate Notch signaling activity, we crossed a Notch signaling reporter mouse CBF:H2B-Venus (27) with two established mouse models of pancreatic cancer, KC (Ptf1a^Cre/+^; Kras^LSL-G12D/+^) and KPC, to generate KC;CBF:H2B-Venus and KPC;CBF:H2B-Venus mice. These models express the Venus fluorescent protein upon Notch signaling activation (Figure 2A and S2A). In the normal pancreas, Venus signal was detected in ductal epithelial cells and was co-localized with HES1, cleaved NOTCH1, and NOTCH2 by immunofluorescent staining (Figure S2B). At very early stages of pancreatic tumorigenesis in 6-week-old KC pancreata, where acinar-ductal metaplasia (ADM) predominates, we observed Venus-expressing macrophages and fibroblasts in the stroma (Figure 2B) by co-immunofluorescent staining of GFP (to detect Venus on FFPE sections) and F4/80 or SMA. Later in 8- and 12-week-old KC and 11-week-old KPC mice we found Pancreatic Intraepithelial Neoplasia (PanIN), a precursor for invasive pancreatic cancer, and more macrophages and fibroblasts in the PanIN microenvironment expressed Venus signifying active Notch signaling (Figure 2C and S2C). Similarly, co-immunofluorescent staining showed Venus-expressing macrophages and fibroblasts in mouse PDA in a 21-week-old KPC mouse (Figure 2D). Based on these observations, Notch signaling activation in the TME starts at the earliest stages of carcinogenesis and persists through disease initiation and progression.

**Figure 2.**
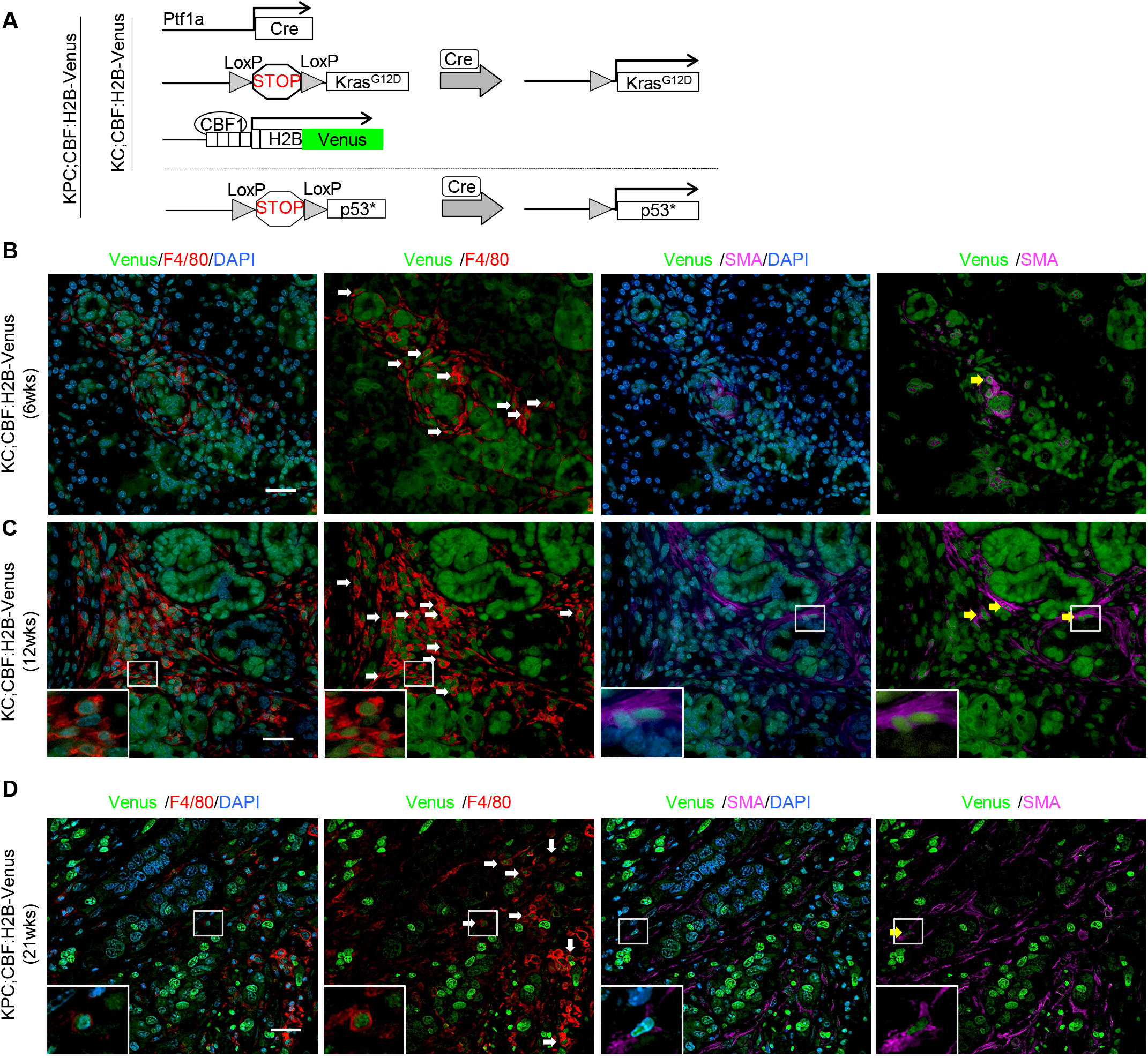
Notch activation detected in the tumor microenvironment of mouse spontaneous PanIN and PDA. (A) Genetic makeup of the KC; CBF:H2B-Venus and KPC; CBF:H2B-Venus mice. (B-D) Co-immunofluorescent staining for Venus (green), F4/80 (red), SMA (magenta) and DAPI (blue) in pancreata harvested from KC; CBF:H2B-Venus and KPC; CBF:H2B-Venus mice at indicated age. Scale bar 50 μm. White arrows show Venus expression in F4/80 positive cells and yellow arrows show Venus expression in SMA positive fibroblasts.

To expand our analysis to full-blown cancer, we orthotopically implanted two syngeneic mouse PDA cell lines, 7940B and MT3-2D (both derived from Pdx1-Cre; Kras^LSL-G12D/+^; Trp53^LSL-R172H/+^ tumors), into C57BL/6J:CBF:H2B-Venus mice and harvested the resulting tumors for flow cytometry analysis after three-weeks (Figure 3A). Venus expression was detected in multiple cell types from the tumor microenvironment including fibroblasts, macrophages, B cells, and T cells (Figure 3B-E). Macrophages comprised up to ~60% of all immune cells (Figure 3C) and about half of the TAMs were Venus^+^, making them the largest Venus-expressing cell population within the TME (Figure 3C and 3F). In contrast, fibroblasts contributed ~20% of total cells and Venus-expressing fibroblasts were rare (Figure 3F and S3A). Corresponding spleen tissues from the tumor bearing mice were used as an internal control and Venus expression was also found in splenic macrophages, B cells and T cells (Figure S3B). To validate that Venus expression represents active Notch signaling we performed co-immunofluorescent staining on the orthotopic PDA sections and determined that F4/80^+^GFP^+^ macrophages were also HES1 positive (Figure S3C). Thus, Notch pathway component and target gene expression characterizes macrophages infiltrating precursor lesions and advanced pancreatic cancer, and Notch activity is reflected by expression of the Venus reporter.

**Figure 3.**
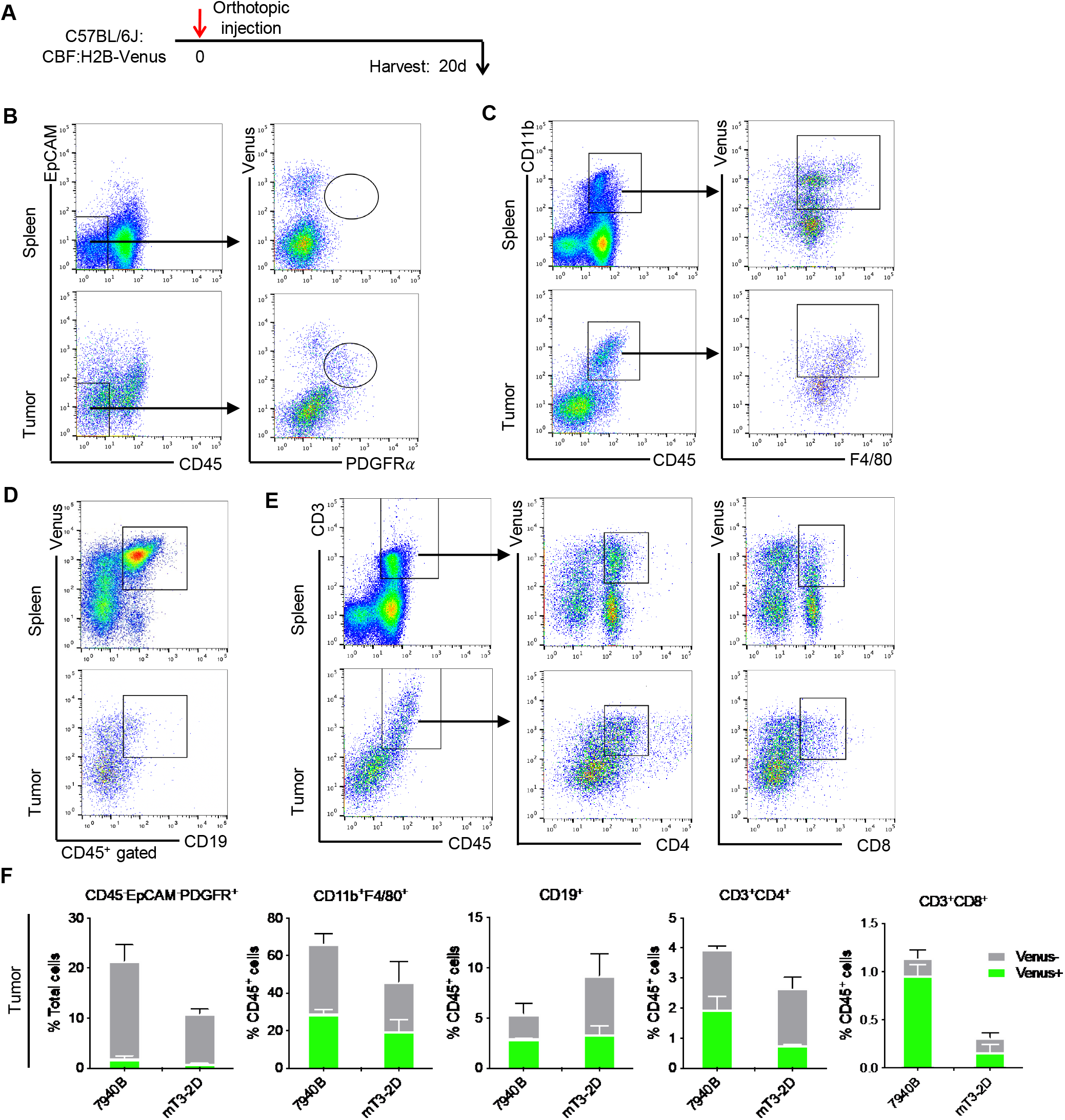
Notch signaling activation in multiple stromal cell types in mouse pancreatic orthotopic tumor. (A) Experimental design. (B) Representative dot plots of flow cytometry analysis of Venus expression in CD45^-^EpCAM^-^PDGFR*α*^+^ fibroblasts, (C) in CD45^+^CD11b^+^F4/80 ^+^macrophages, (D) in CD45^+^D19^+^ B cells, (E) in CD45^+^CD3^+^CD4^+^ or CD8^+^ T cells derived from spleens or pancreatic tumors harvested from PDA bearing mice. (F) Venus positive or negative fibroblasts (CD45^-^EpCAM^-^PDGFR*α*^+^), macrophages (CD45^+^CD11b^+^F4/80^+^), B (CD45^+^CD19^+^), CD4 T (CD45^+^CD3^+^CD4^+^) and CD8 T (CD45^+^CD3^+^CD8^+^) cells in pancreatic tumors were measured by flow cytometry. Data represent mean ± SEM.

### Notch signaling in tumor-associated macrophages correlates with immunosuppressive polarization

To functionally characterize the role of Notch signaling in tumor associated macrophages, we used fluorescence-activated cell sorting to isolate Venus^+^ and Venus^-^ TAMs from the orthotopic 7940B tumors. The Venus^+^ and Venus^-^ TAM populations were similar in cell number (Figure 4A), consistent with co-immunofluorescent staining showing about half of the F4/80^+^ macrophages expressing Venus (Figure 4B). We then performed qRT-PCR to molecularly characterize them (Figure 4C). First, we confirmed that *Hes1* expression, a measure of Notch signaling, was higher in Venus^+^ TAMs compared to Venus^-^ TAMs. Interestingly, we also found that Wnt signaling target *Lef1* and several alternative activated macrophage (M2-like) markers, including *Arg1*, *Msr1* and *Mrc1*, were elevated in Venus^+^ TAMs (Figure 4C). Consistent with these findings, expression of immunosuppressive cytokines and chemokines such as *Il10* and *Tgfβ1* was also higher in Venus^+^ TAMs. Finally, the classically activated macrophage (M1) marker *Nos2* was lower in Venus^+^ TAMs and there were no significant differences in expression levels of *Notch1*, *Axin2*, *Tnfα* and *Chi3l3* between the two TAM subsets. Overall, gene expression analysis pointed to a positive correlation between Notch activation and M2-like polarization of TAMs.

**Figure 4.**
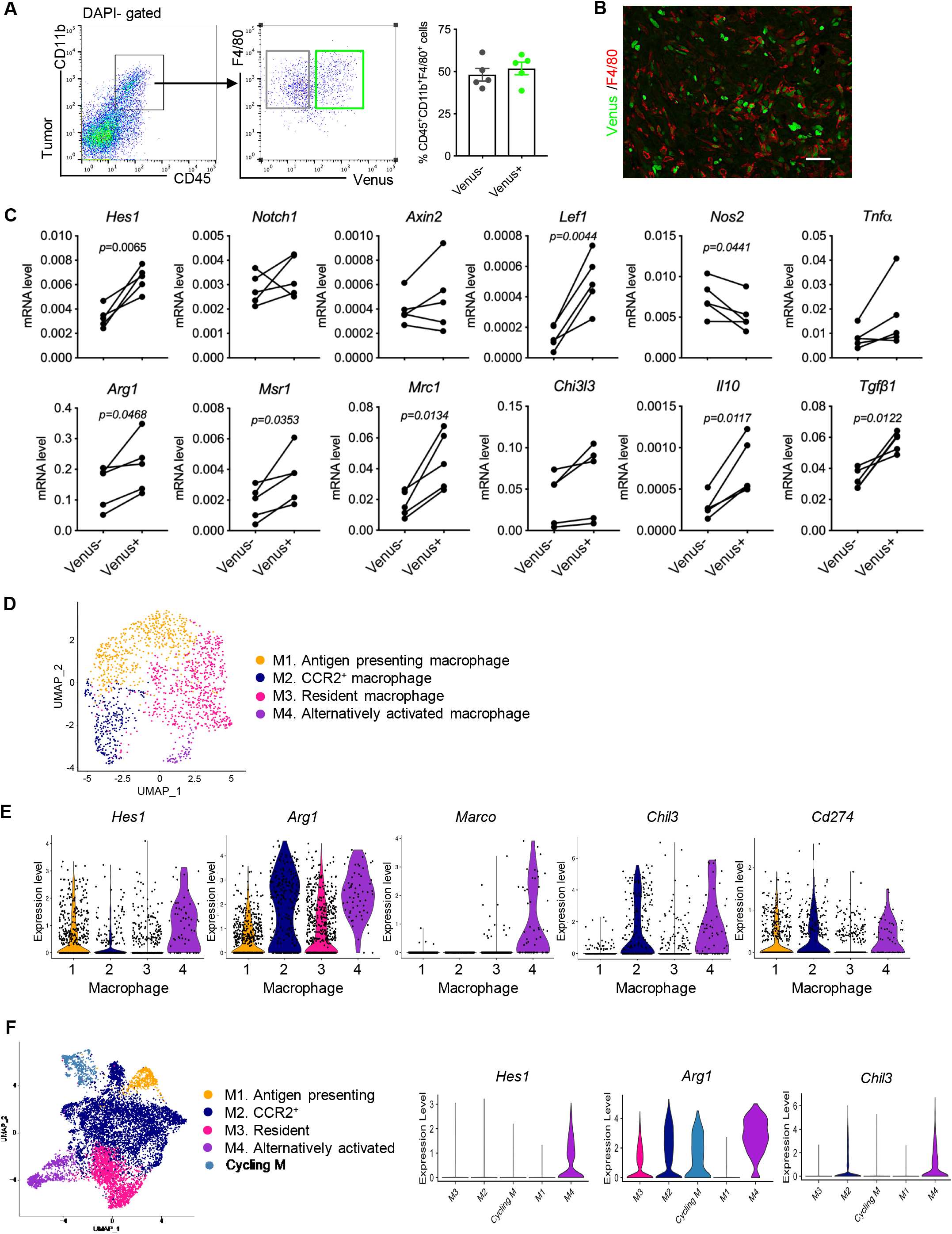
Notch signaling activation is prevalent in alternatively activated TAMs. (A) Fluorescence-activated cell sorting for Venus positive or negative tumor associated macrophages. Data represent mean ± SEM. (B) Co-immunofluorescent staining for Venus (green), and F4/80 (red) in pancreata harvested from orthotopic PDA. Scale bar 50 μm. (C) qRT-PCR for *Hes1*, *Notch1*, *Axin2*, *Lef1*, *Nos2*, *Tnfα*, *Arg1*, *Msr1*, *Mrc1*, *Chi3l3*, *Il10* and *Tgfβ1* expression in Venus negative or positive tumor associated macrophages. Data represent mean ± SEM, n=5. The statistical difference was determined by two-tailed t-tests. (D) UMAP plot of four macrophage sub-populations identified from single-cell RNA sequencing analysis with KPC mouse pancreatic cancer, color-coded by their associated cluster. (E) Violin plots of single cell RNA sequencing analysis showing expression levels of *Hes1*, *Arg1*, *Marco*, *Chil3* and *Cd274* in different macrophage subsets derived from mouse pancreatic cancer sample. (F) UMAP plot of four macrophage sub-populations identified from a second single-cell RNA sequencing dataset (E Elyada, 2019 dataset) of KPC mouse pancreatic cancer, and Violin plots of expression of *Hes1*, *Arg1* and *Chil3*.

To validate our findings, we further analyzed the single-cell sequencing data obtained from the KPC model. We identified four subsets of macrophages (Figure 4D and S4A) and performed gene expression analysis for Notch pathway genes (Figure S4B). All four TAM subsets expressed Notch receptors at varying levels with highest expression noted for *Notch1* and *Notch2*. The macrophage 4 population had the highest expression level of *Hes1*, indicating high levels of Notch signaling. The same macrophage 4 population also expressed several M2/tumor associated macrophage markers including *Arg1, Marco* and *Chi3l3* and had the highest level of the immune checkpoint ligand *Cd274 (PD-L1)* (Figure 4E). We further analyzed single-cell RNAseq data from a publicly available KPC (Pdx1-Cre; Kras^LSL-G12D/+^; Trp53^LSL-R172H/+^) tumor dataset (28). In this case, the *Hes1* high population also expressed the highest levels of *Arg1* and *Chi3l3* (Figure 4F and Figure S4C).

To verify our findings in human PDA, we isolated myeloid cells including macrophages and granulocytes from our previously published human scRNA-seq immune dataset for an in-depth analysis (25). As in the mouse, *HES1* was expressed in myeloid cells, and its expression is more abundant in PDA compared to adjacent and normal pancreata (Figure 5A and 5B). Among the four identified TAM subsets in human PDA, CCR2^+^ classical macrophages, resident macrophages and alternatively activated macrophages all had similar amount of *HES1* expression, which in turn was higher compared to other myeloid subsets (Figure 5C). To seek a better understanding of the role of Notch signaling activation in TAMs we compared the gene expression profile between *HES1*-positive and *HES1*-negative macrophages from human patients. We discovered higher expression of complement genes *C1QA, C1QB*, alternatively activated macrophage markers *CD163*, *MSR1*, *MRC1* and *MARCO*, and resident macrophage marker *STAB1* in *HES1*-positive macrophages. We previously identified complement genes *C1QA*, *C1QB*, and *TREM2* as part of a pancreatic tumor-specific signature in macrophages (29). *HES1*-positive macrophages also have higher levels of *HIF1A*, potentially linking Notch signaling activation in macrophages to hypoxia in the TME (Figure 5D).

**Figure 5.**
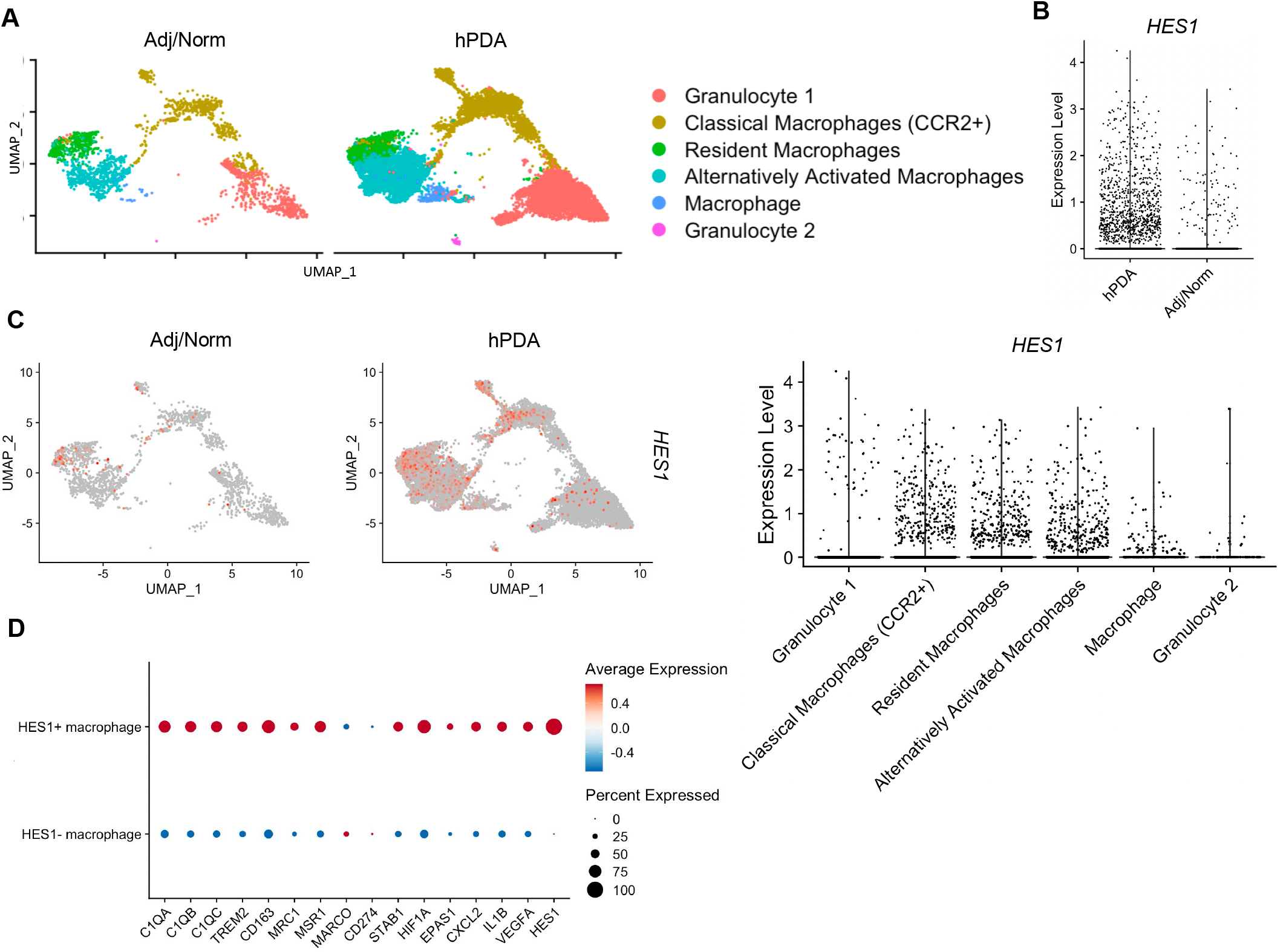
Notch signaling activation in human TAMs correlates with tumor-specific gene signature. (A) UMAP plot showing six myeloid cell sub-populations identified from single-cell RNA sequencing analysis with human pancreatic cancer samples (n=16) and adjacent benign/normal tissues (n=3), color-coded by their associated cluster. (B) Violin plot of single cell RNA sequencing analysis comparing expression level of *HES1* in myeloid cells between human pancreatic cancer samples and adjacent benign/normal tissues. (C) Feature plot and Violin plot of single cell RNA sequencing analysis showing expression level of *HES1* in myeloid cells derived from human pancreatic cancer samples and adjacent benign/normal tissues. (D) Dot plots of single cell RNA sequencing analysis showing differentially expressed genes of *C1QA*, *C1QB*, *CD163*, *MSR1*, *MRC1*, *STAB1* and *HIF1A* between *HES1*-positive and *HES1*-negative macrophage subsets derived from human pancreatic samples.

To elucidate a potential causal link between Notch activation and immunosuppressive function of TAMs, we developed an *in vitro* co-culture system in which bone marrow derived myeloid cells (BMDM) are differentiated and polarized to TAMs by cancer conditioned media or direct co-culture with PDA cells (Figure 6A). Using BMDM derived from C57BL/6J:CBF:H2B-Venus mice we traced activation of Notch signaling by Venus expression. We observed Venus expression only in BMDM cells co-cultured with PDA cells directly and not with cancer conditioned media only, supporting the notion that direct physical contact between the tumor cells and myeloid cells is needed for Notch activation as expected (Figure 6B). The Venus expressing BMDM cells also expressed F4/80 and Arg1, markers of immunosuppressive macrophages (Figure 6C). Moreover, when BMDM co-cultured with PDA cells were treated with the γ-secretase inhibitor Crenigacestat (30) to inhibit the proteolytic signaling downstream of Notch receptors, the expression of *Hes1*, *Arg1*, *Chi3l3*, *Mrc1*, *Tgfβ1* or *Il10* were down-regulated compared to vehicle-treated cells (Figure 6D and 6E). These results point to direct tumor epithelial/macrophage contact in the PDA microenvironment activating macrophage Notch signaling and resulting in polarization of TAMs to an immunosuppressive phenotype.

**Figure 6.**
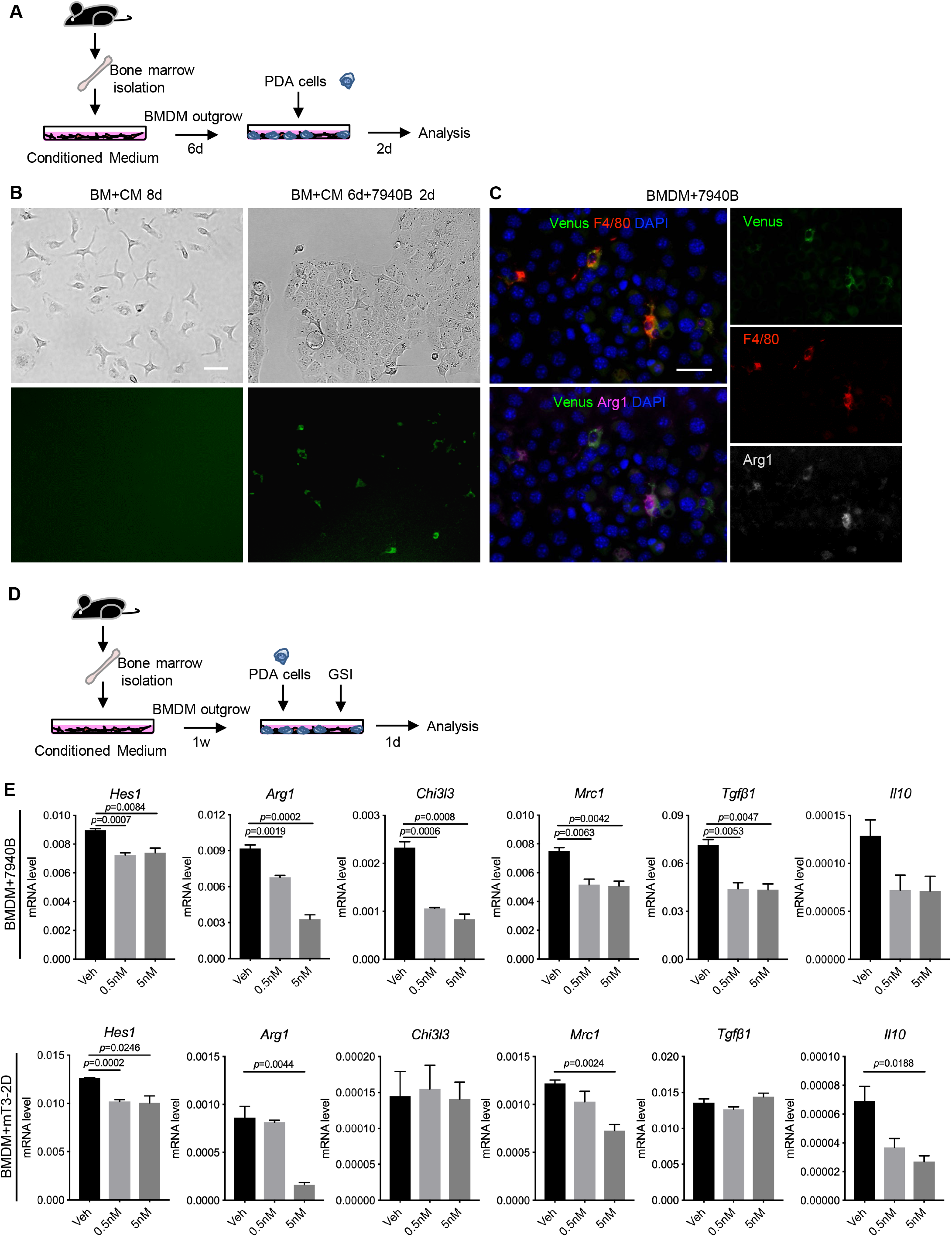
γ-secretase inhibitor reduced immunosuppressive markers expression in TAM. (A) Experimental design of bone marrow derived macrophage (BMDM) co-cultured with PDA cancer cells. (B) Bright field and fluorescent microscopy images of bone marrow (BM) cells in culture with cancer cell conditioned medium (CM) or cancer cells 7940B. (C) Venus (green) and co-immunofluorescent staining for F4/80 (red) and Arg1 (magenta) in BMDM co-cultured with 7940B cancer cells. Scale bar 50 μm. (C) Experimental design of bone marrow derived macrophage (BMDM) co-cultured with PDA cancer cells and treated with γ-secretase inhibitor (GSI). (D) qRT-PCR for *Hes1*, *Arg1*, *Chi3l3*, *Mrc1*, *Tgfβ1* and *Il10* expression in vehicle or GSI (Crenigacestat at 0.5 nM or 5 nM) treated BMDM co-cultured with PDA cells 7940B and mT32D. Data represent mean ± SEM, n=3. The statistical difference was determined by two-tailed t-tests.

### Inhibition of Notch signaling synergizes with PD1 blockade to activate anti-tumor immune activity

Given our functional results implicating epithelial/macrophage crosstalk in Notch activation and immunosuppressive TAM polarization, we sought to investigate the therapeutic and immunomodulatory potential of Notch signaling inhibition *in vivo*. We implanted 7940B cells orthotopically into syngeneic C57BL/6J mice; a week later, tumor bearing mice were treated with either GSI, anti-PD1, or combination of both (Figure 7A). At harvest (day 19) we observed smaller tumors in the combination treatment cohort compared to the controls or either treatment alone (Figure 7B). Histology analysis of the orthotopic tumors revealed an increase in tumor-infiltrating CD8^+^ T cells along with increased levels of the T cell activation marker granzyme B in the combination treatment cohort (Figure 7C and quantification in Figure 7E). We also observed increased cell apoptosis by cleaved caspase 3 immunohistochemistry staining after GSI and anti-PD1 combination treatment (Figure 7D and quantification in Figure 7F). Targeting Notch signaling alone (GSI group) also showed a trend in increase of tumor infiltrating CD8 T cells although there was no change in Granzyme B production or cell apoptosis compared to the control group. In contrast, the tumor infiltrating macrophage number remained unchanged in all conditions (Figure S5A and quantification in Figure S5B). These findings are consistent with activation of a productive anti-tumor immune response upon combined treatment targeting Notch signaling and the PD1 checkpoint. The results also support the notion that native Notch signaling promotes immunosuppressive TAM polarization and targeting Notch may be a useful addition to immunotherapy approaches in pancreatic cancer.

**Figure 7.**
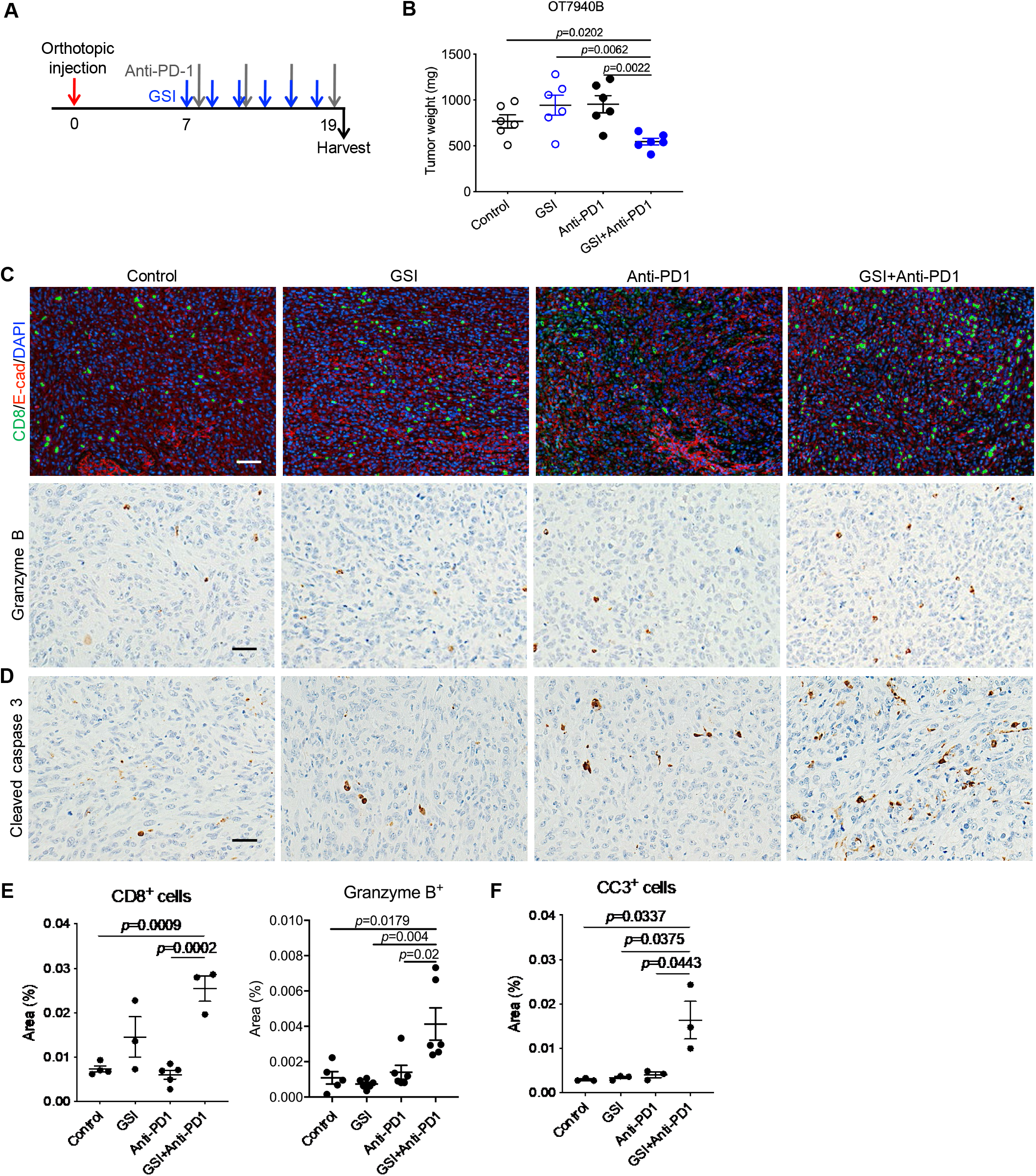
Inhibition of Notch signaling combined with PD1 blockade increases anti-tumor immune activity. (A) Experimental design of orthotopic implantation of pancreatic cancer cells 7940B. (B) Tumor weights of orthotopic PDA harvested from mice that received vehicle/IgG as control, or GSI, or anti-PD1 or combination of GSI and anti-PD1. Data represent mean ± SEM, n=6. The statistical difference was determined by two-tailed t-tests. (C) Co-immunofluorescent staining for CD8 (green), E-cad (red) and DAPI (blue), and immunohistochemical staining for Granzyme B in orthotopic PDA tumors. Scale bar 50 μm. (D) Quantification of CD8, Granzyme B positive area (%). Data represent mean ± SEM, n=3~6. The statistical difference was determined by two-tailed t-tests. (E) Immunohistochemical staining for Cleaved caspase 3 in orthotopic PDA tumors. Scale bar 50 μm. (F) Quantification of Cleaved caspase 3 (CC3) positive area (%). Data represent mean ± SEM, n=3~6. The statistical difference was determined by two-tailed t-tests.

## Discussion

Immunotherapy has so far not been effective as a treatment option for pancreatic cancer (31,32). Specifically, targeting the PD1 immune checkpoint with a single agent provides no benefit to patients; effective combination approaches are thus needed. The complex immunosuppressive nature of the PDA tumor microenvironment is the direct cause of this and reflects the interplay of many different cell types (7). The tumor-associated macrophages play a key immunosuppressive role within this context in pancreatic cancer (33–36).

Notch signaling has been previously implicated in myeloid development in the bone marrow as well as in local functional maturation in the tissue/tumor microenvironment (19,37). In addition, tumor epithelial Notch signaling alters the tumor secretome to promote an immunosuppressive response in the surrounding microenvironment (38). Role for direct Notch signaling in myeloid cells has been lacking in PDA despite evidence for it in other tumor types (18). Our paper demonstrates that Notch pathway components and active signaling are present in multiple cell populations in both human and murine tumors including the macrophages, T cells, fibroblasts, and the endothelium. We also use a non-invasive Notch pathway activity reporter to identify and characterize the phenotype and function of the myeloid cells in pancreatic tumors. Our data support the notion that Notch-active myeloid cells are primarily tumor-associated macrophages with a more immunosuppressive M2-like/alternatively activated phenotype marked by higher expression of immunosuppressive cytokines, arginase 1, and immune checkpoint molecules.

Contrary to our results, other investigators have found that Notch activation primarily drives an M1-like, tumor suppressive phenotype in macrophages (22,39). In addition, independent modulation of Notch signaling via Notch intracellular domain overexpression or deletion of Rbpj – a key transcriptional transducer of Notch signaling – in the myeloid compartment of an autochthonous mouse model of pancreatic neoplasia suggested that activation of Notch signaling in myeloid cells drives a stronger anti-tumor immune response (23). The discrepancy between these findings and our observations in myeloid cells with intact, native level of signaling is potentially at least partially explained by the modular and combinatorial nature of the Notch signaling pathway (40–44). Final Notch signaling response is a product of integration of both cis and trans cellular signaling and inhibition. In addition to this, different Notch ligands can evoke distinct quantitative and qualitative cellular responses (44). Work published prior to our manuscript often relied on high level of activation or inhibition of Notch signaling through isolated ligand or Notch intracellular domain overexpression or complete genetic abrogation of the Notch transcriptional response. In addition, many of these interventions were present throughout myeloid development, migration, and local microenvironment differentiation, which may lead to signaling and functional myeloid states that would not normally be seen in an intact in vivo context. In comparison, our work utilized a non-invasive functional fluorescent reporter of Notch activity to identify cells with native levels of Notch signaling activation.

This allowed us to isolate and molecularly and functionally characterize the myeloid cells with a productive Notch transcriptional response without disrupting any of the native activating and inhibitory receptor/ligand interactions. The immunosuppressive nature of these myeloid cells could be partially reversed by γ-secretase inhibition, which blocks the proteolytic activation of Notch signaling. As a final functional proof of concept, we demonstrated that combining γ-secretase inhibition with immune checkpoint inhibition leads to a productive anti-tumor immune response with increased infiltration of CD8^+^ cytotoxic T cells and increased tumor apoptosis. Overall, our data shows that the effects of Notch signaling in the PDA tumor microenvironment are highly ligand, receptor, and overall context-dependent.

In addition to demonstrating direct Notch activation and function in the myeloid compartment, our work also implicates Notch in multiple other cell types in the PDA tumor microenvironment including T cells, fibroblasts, and the endothelium. Role of Notch in T cell maturation and function is well documented, but is less clear in the specific context of PDA (45,46). Notch signaling has also been shown to regulate fibroblast phenotype and function in the tumor microenvironment and other inflammatory contexts (47,48). In PDA, cancer-associated fibroblasts can be parsed into multiple coexisting subtypes regulating the tumor microenvironment and immune response (28,49–57). We and others have previously shown that dysregulation of Hedgehog signaling plays a key role in CAF function and immune polarization within the TME (49,51,58–60). How Notch integrates with Hedgehog and other signaling pathways in the CAFs and how this affects the tumor microenvironment remains an active area of study. Similarly, Notch ligands are also highly expressed on tumor-associated endothelium (47). Myeloid cells recruited to the tumor microenvironment have to traverse the endothelium as the first barrier during their extravasation into the tumor and endothelial Notch ligands may serve as the first relevant Notch signal that leading to myeloid polarization in the TME. Along these lines radiation-induced increased expression of the Notch ligand Jagged1 in lung endothelium leads to alternative M2-like polarization of myeloid cells as they are recruited into the lung parenchyma (61). Similar mechanisms are potentially at play in the PDA TME. In summary our work implicates direct Notch signaling in immunosuppressive myeloid response polarization in pancreatic cancer and suggests the presence of a complex Notch-specific network of interactions between various cell types regulating this polarization. Further understanding of Notch signaling and its role in pancreatic cancer may provide future alternative means to redeploy already-developed Notch-modulating drugs in combination chemoimmunotherapy regimens.

## Acknowledgements

We thank Dr. Howard Crawford and Daniel Long for histological services. This project was also supported by the Tissue and Molecular Pathology and Flow Cytometry Shared Resources at the Rogel Cancer Center and the University of Michigan Advanced Genomics Core.

