## Supplementary data and information for "Notch signaling regulates immunosuppressive tumor-associated macrophage function in pancreatic cancer"

**Supplemental Figure 1. Notch activation in the tumor microenvironment of human PDA.**

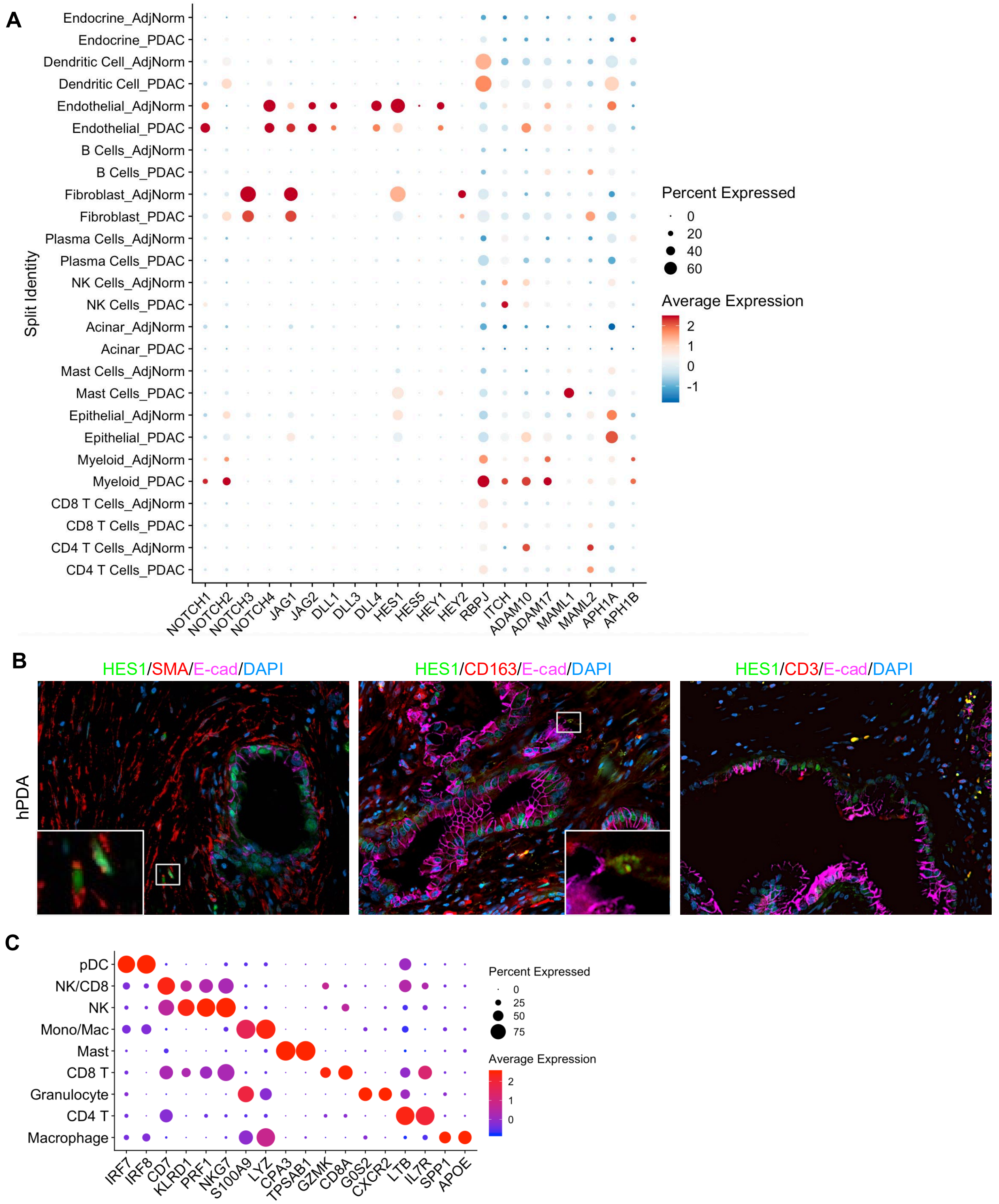

**Supplemental Figure 1. Notch activation in the tumor microenvironment of human PDA.**

(A) Dot plots showing expression of Notch pathway genes across all cell clusters identified in the single-cell RNA sequencing analysis of human pancreatic samples. Size of dots represents percentage of cells expressing a particular gene and intensity of color indicates level of mean expression. (B) Co-immunofluorescent staining for HES1 (green), SMA, CD163 or CD3 (red), E-cad (magenta) and DAPI (blue) in human PDA sample. Scale bar 50 μm. (C) Dot plots showing lineage markers for immune cell clusters identified from human single-cell RNA sequencing samples.

**Supplemental Figure 2. Notch activation detected in C57BL/6J:CBF:H2B-Venus mouse pancreas and tumor microenvironment of spontaneous PanIN lesions.**

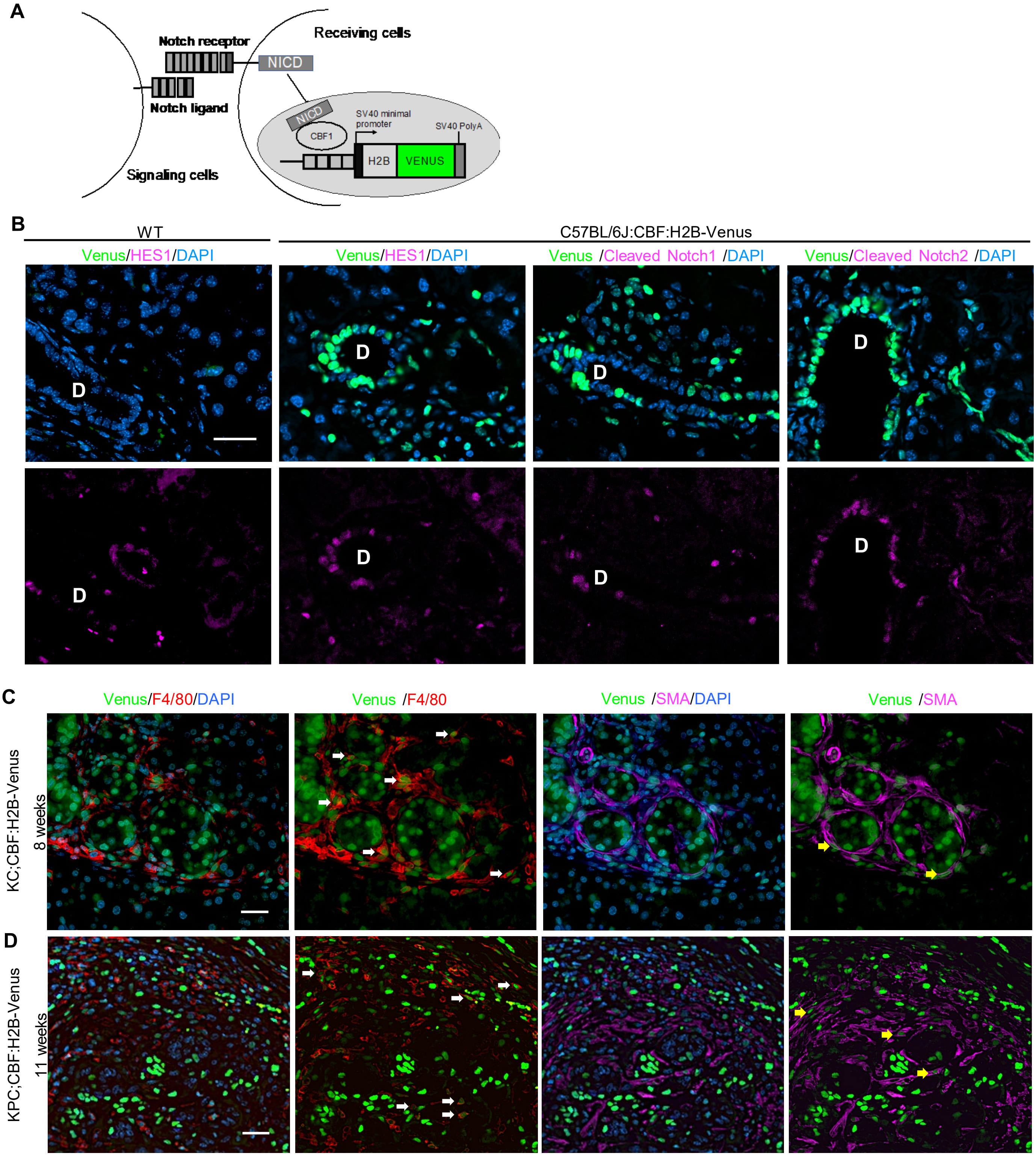

**Supplemental Figure 2. Notch activation detected in normal pancreas of C57BL/6J:CBF:H2B-Venus mouse and tumor microenvironment of spontaneous PanIN lesions.**

(A) *C57BL/6J:CBF:H2B-Venus* mouse model. The transgenic H2B-Venus reporter construct contains multiple Notch responsive binding sites. When the Notch pathway is active in the cells, the H2B-Venus fluorescent protein reporter is expressed in the cell's nucleus. (B) Immunofluorescent staining of Hes1, cleaved Notach1 or 2 in the normal pancreas from the wild type (WT) and *C57BL/6J:CBF:H2B-Venus* mouse. D: duct. Scale bar 50 µm. (C) Co-immunofluorescent staining for Venus (green), F4/80 (red), SMA (magenta) and DAPI (blue) in pancreata harvested from KC; CBF:H2B-Venus and (D) KPC; CBF:H2B-Venus mice at indicated age. Scale bar 50 µm. White arrows show Venus expression in F4/80 positive cells and yellow arrows show Venus expression in SMA positive fibroblasts.

**Supplemental Figure 3. Notch activation detected in C57BL/6J:CBF:H2B-Venus mouse pancreatic orthotopic tumor.**

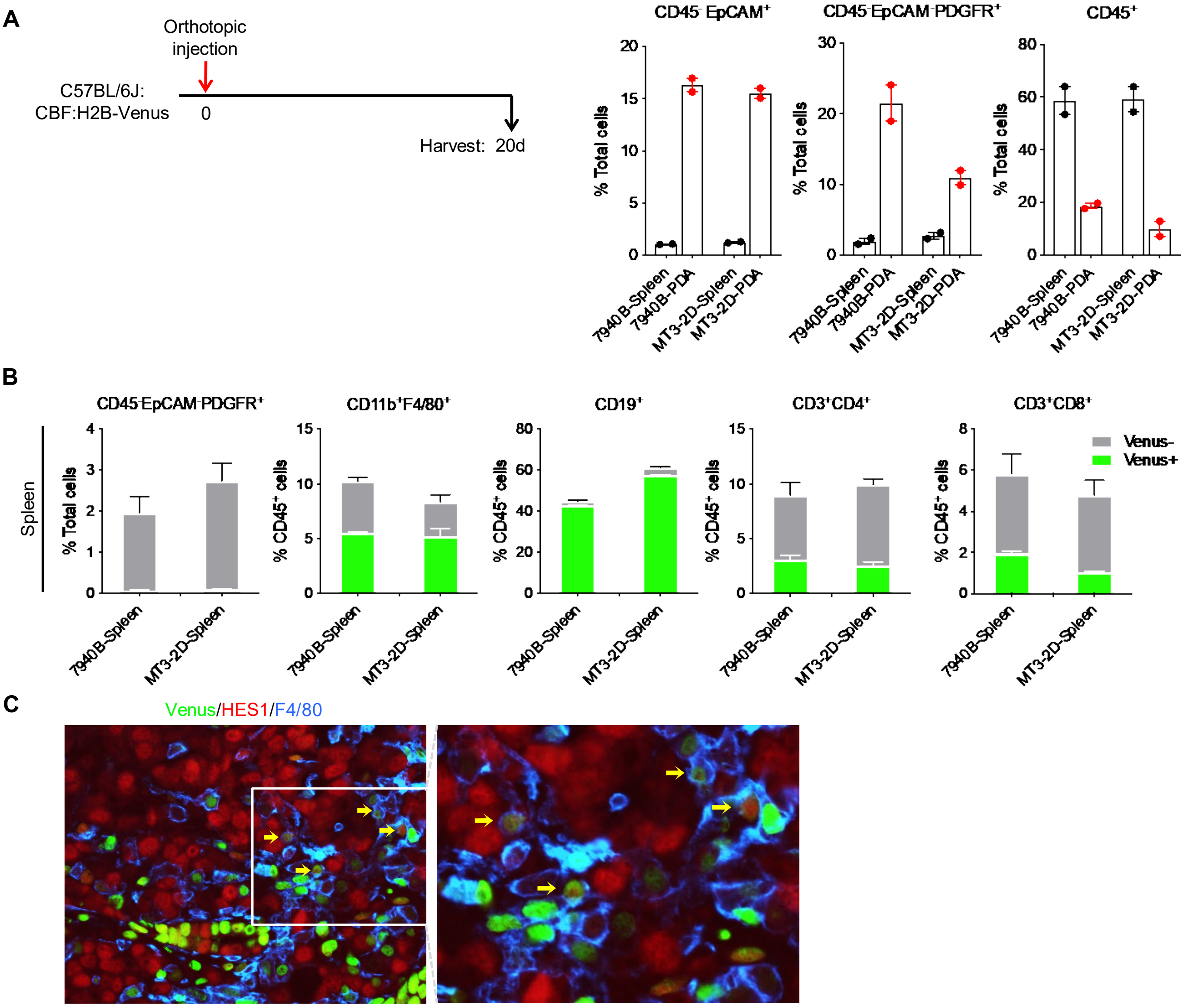

**Supplemental Figure 3. Notch activation detected in C57BL/6J:CBF:H2B-Venus mouse pancreatic orthotopic tumor.**

(A) Percentages of CD45-EpCAM+ tumor epithelial cells, CD45-EpCAM-PDGFR$\alpha$+ fibroblasts, and CD45+ immune cells in spleen or pancreata harvested from PDA bearing mice were measured by flow cytometry. Data represent mean ± SEM. (B) Venus positive or negative fibroblasts (CD45$^{-}$EpCAM$^{-}$PDGFR$\alpha^{+}$), macrophages (CD45$^{+}$CD11b$^{+}$F4/80$^{+}$), B (CD45$^{+}$CD19$^{+}$), CD4 T (CD45$^{+}$CD3$^{+}$CD4$^{+}$) and CD8 T (CD45$^{+}$CD3$^{+}$CD8$^{+}$) cells in spleens from PDA bearing mice were measured by flow cytometry. Data represent mean ± SEM. (C) Co-immunofluorescent staining for Venus (green), HES1 (red), F4/80 (blue) in mouse pancreatic orthotopic tumor sample. Scale bar 50 µm.

**Supplemental Figure 4. Single cell analysis of Notch signaling activation in TAMs.**

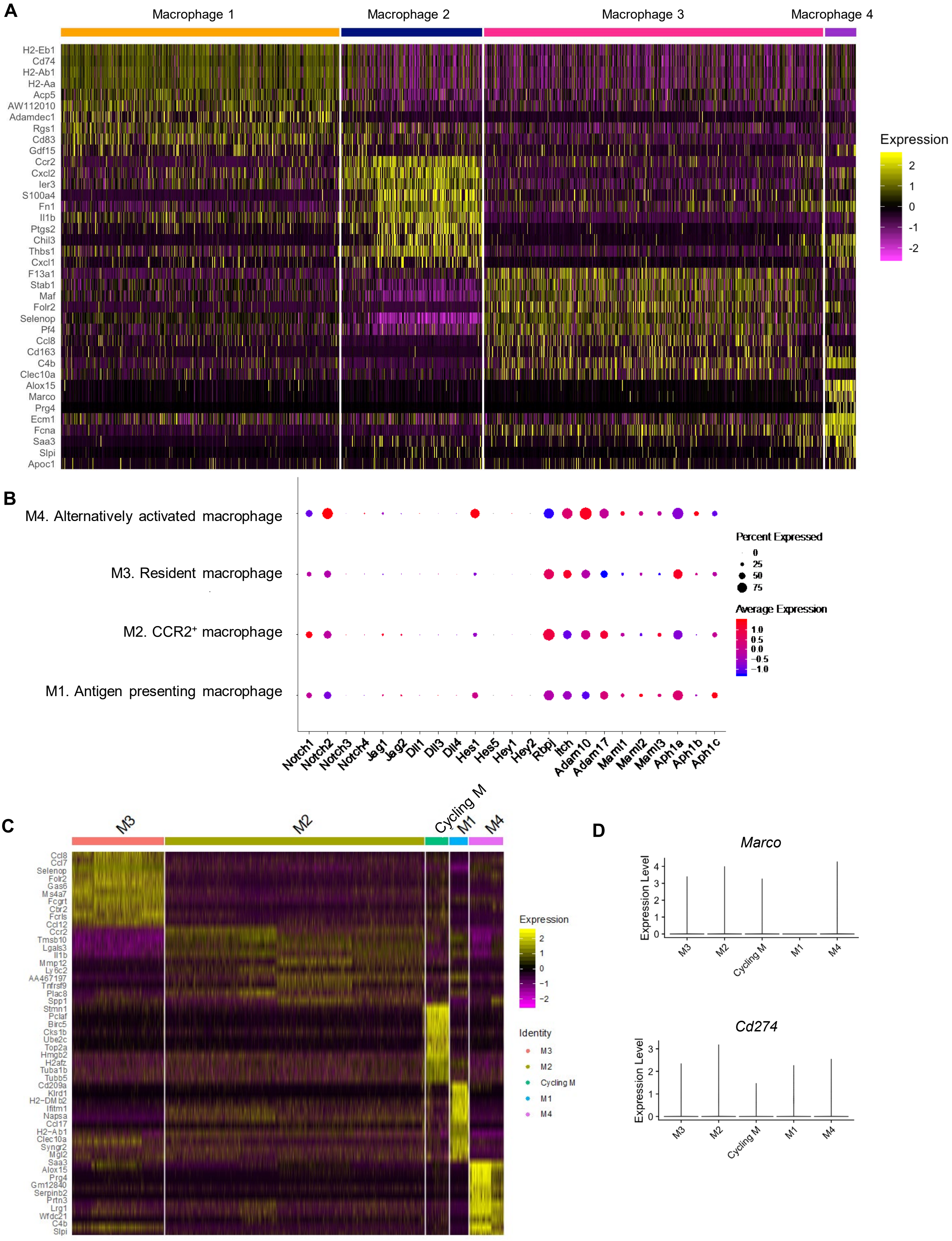

**Supplemental Figure 4. Single cell analysis of Notch signaling activation in TAMs.**

(A) Heatmap showing the top most differentially expressed genes in the four macrophage subsets identified from single-cell RNA sequencing analysis with KPC mouse pancreatic cancer. (B) Dot plots showing expression of Notch pathway genes across the four macrophage subsets identified in the single-cell RNA sequencing analysis with KPC mouse pancreatic cancer. Size of dots represents percentage of cells expressing a particular gene and intensity of color indicates level of mean expression. (C) Heatmap showing the top most differentially expressed genes in the five macrophage subsets identified from the second set of single-cell RNA sequencing analysis with KPC mouse pancreatic cancer (28). (D) Violin plots showing *Marco* and *Cd274* expression in KPC macrophages (28).

**Supplemental Figure 5. γ-secretase inhibitor treatment reduces expression of HES1 *in vivo*.**

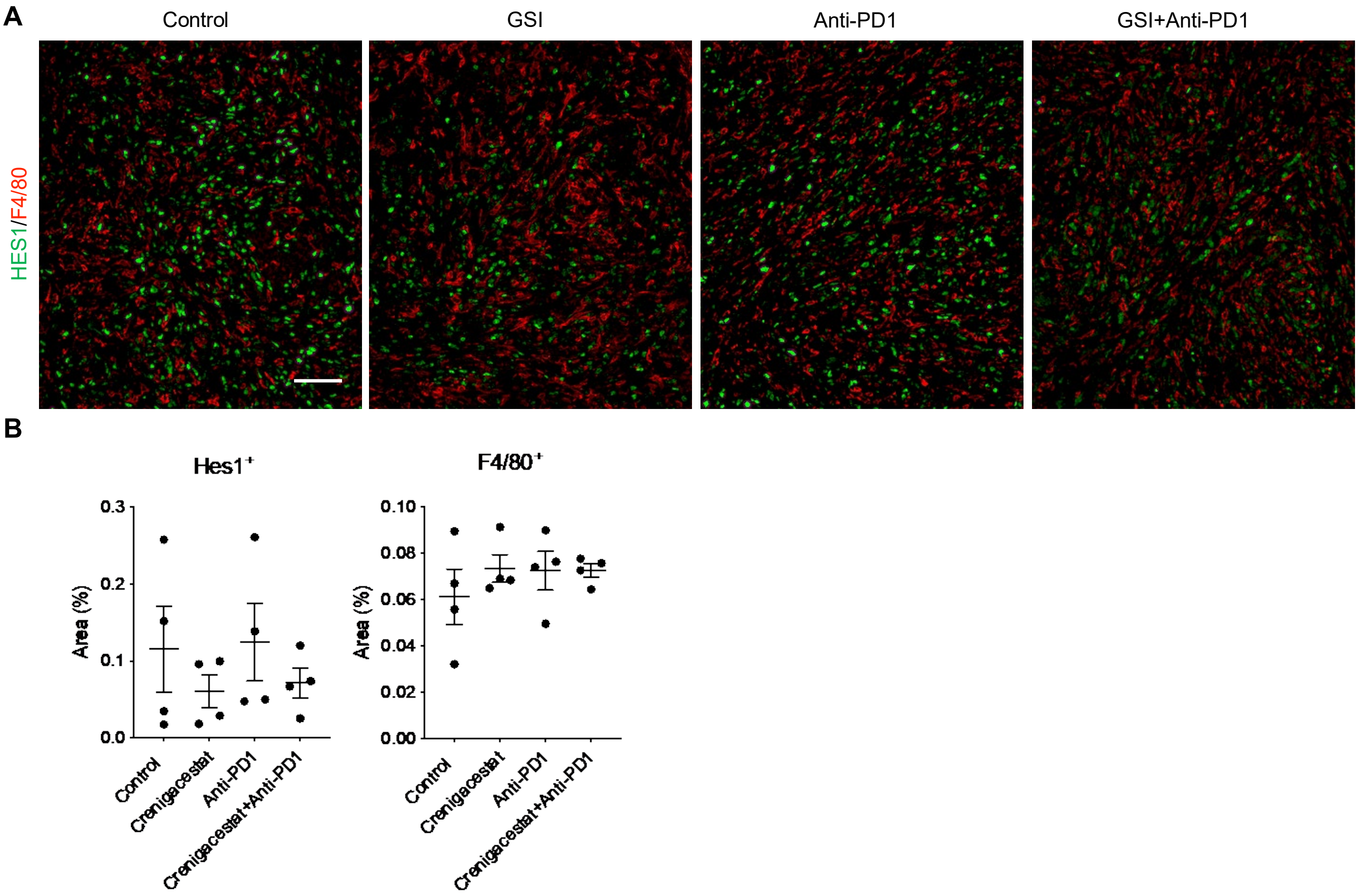

**Supplemental Figure 5. γ-secretase inhibitor treatment reduces expression of HES1 *in vivo*.**

(A) Co-immunofluorescent staining for HES1 (green) and F4/80 (red) in orthotopic PDA tumors. Scale bar 50 μm. (B) Quantification of HES1 and F4/80 positive area (%). Data represent mean ± SEM, n=4.

**Supplemental Table 1: Antibodies**

| Antibody | Supplier | Catalog Number | IHC dilution | IF dilution |
|---|---|---|---|---|
| Arginase-1 | Cell Signaling | 93668 | - | 1:100 |
| Alpha-smooth muscle actin | Sigma-Aldrich | A2547 | - | 1:1000 |
| CD3 | Dako | A0452 | - | 1:200 |
| CD8 | Cell Signaling | 98941 | - | 1:400 |
| CD31 | Invitrogen | PA5-16301 | - | 1:50 |
| CD163 | Cell Signaling | 93498 | - | 1:150 |
| CK19 (TromaIII) | Iowa Developmental Hybridoma Bank | - | - | 1:50 |
| Cleaved Caspase 3 | Cell Signaling | 9661 | 1:300 | - |
| Cleaved Notch1 | Cell Signaling | 4147 | - | 1:200 |
| Cleaved Notch2 | Abcam | ab52302 | - | 1:100 |
| E-cadherin (4A2) | Cell Signaling | 14472 | - | 1:50 |
| E-cadherin (24E10) | Cell Signaling | 3195 | - | 1:100 |
| F4/80 | Cell Signaling | 70076 | - | 1:200 |
| GFP | Abcam | Ab6673 | - | 1:200 |
| Granzyme B | Cell Signaling | 17215 | 1:200 | - |
| HES1 | Cell Signaling | 11988 | - | 1:200 |
| **Flow Cytometry Antibody** | **Supplier** | **Clone** | **Dilution** | |
| CD3 | BD Pharmingen | 17A2 | 1:100 | |
| CD11b | BD Pharmingen | M1/70 | 1:100 | |
| CD19 | BD Pharmingen | 1D3 | 1:100 | |
| CD45 | BD Bioscience | 30-F11 | 1:100 | |
| CD326 (EpCAM) | BioLegend | G8.8 | 1:100 | |
| F4/80 | eBioscience | BM8 | 1:100 | |

| | | | |
|---|---|---|---|
| PDGFRa | eBioscience | APA5 | 1:100 |

**Supplemental Table 2: Primer sequences for quantitative RT-PCR**

| Gene | Forward Primer | Reverse Primer |
|---|---|---|
| *Arg1* | CTCCAAGCCAAAGTCCTTAGAG | AGGAGCTGTCATTAGGGACATC |
| *Axin2* | GCCAATGGCCAAGTGTCTCT | GCGTCATCTCCTTGGGCA |
| *Chi3l3* | CAGGTCTGGCAATTCTTCTGAA | GTCTTGCTCATGTGTGTAAGTGA |
| *Cyclophilin A* | TCACAGAATTATTCCAGGATTCATG | TGCCGCCAGTGCCATT |
| *Hes1* | CCAGCCAGTGTCAACACGA | AATGCCGGGAGCTATCTTTCT |
| *Il10* | GCTATGCTGCCTGCTCTTACT | CCTGCTGATCCTCATGCCA |
| *Lef1* | AGTGCAGCTATCAACCAGATCCT | TTTCCGTGCTAGTTCATAGTATTTGG |
| *Mrc1* | CTCTGTTCAGCTATTGGACGC | CGGAATTTCTGGGATTCAGCTTC |
| *Msr1* | GCACAATCTGTGATGATCGCT | CCCAGCATCTTCTGAATGTGAA |
| *Nos2* | GTTCTCAGCCCAACAATACAAGA | GTGGACGGGTCGATGTCAC |
| *Notch1* | ATGCCAGGACTCCAATCCTTG | CGTTCCAGCATTCTTACACGG |
| *Tgfβ1* | TGACGTCACTGGAGTTGTACGG | GGTTCATGTCATGGATGGTGC |
| *Tnfα* | CATCTTCTCAAAATTCGAGTGACAA | TGGGAGTAGACAAGGTACAACCC |
